# Stem cells commit to differentiation following multiple induction events in the Drosophila testis

**DOI:** 10.1101/2021.05.20.444930

**Authors:** Alice C Yuen, Kenzo-Hugo Hillion, Marc Amoyel

**Author notes:** Department of Computational Biology, USR3756 CNRS, Institut Pasteur, 25-28 rue du docteur Roux, 75015 Paris, France.

## Abstract

How and when potential becomes restricted in differentiating stem cell daughters is poorly understood. While it is thought that signals from the niche are actively required to prevent differentiation, another model proposes that stem cells can reversibly transit between multiple states, some of which are primed, but not committed, to differentiate. In the Drosophila testis, somatic cyst stem cells (CySCs) generate cyst cells, which encapsulate the germline to support its development. We find that CySCs are maintained independently of niche self-renewal signals if activity of the PI3K/Tor pathway is inhibited. Conversely, PI3K/Tor is not sufficient alone to drive differentiation, suggesting that it acts to license cells for differentiation. Indeed, we find that the germline is required for differentiation of CySCs in response to PI3K/Tor elevation, indicating that final commitment to differentiation involves several steps and intercellular communication. We propose that CySC daughter cells are plastic, that their fate depends on the availability of neighbouring germ cells, and that PI3K/Tor acts to induce a primed state for CySC daughters to enable coordinated differentiation with the germline.

## Introduction

In tissues with high turnover, stem cells are crucial to maintaining homeostasis by producing new differentiated offspring to replace lost cells. Ensuring long-term tissue maintenance requires that stem cells self-renew, in addition to producing differentiating daughters. Self-renewal capacity was shown early on not to be intrinsic to a stem cell, but to rely instead on the local environment of the cell, termed a niche (Schofield, 1978). The first such niche was identified in the Drosophila ovary (Xie and Spradling, 1998, 2000), and our understanding of stem cell-niche interactions has greatly increased since then (Jones and Wagers, 2008; Morrison and Spradling, 2008). A common feature of many niches is that the signals they produce actively inhibit differentiation of resident stem cells, exemplified by repression of the differentiation factor *bag of marbles (bam)* by niche-derived Bone Morphogenetic Protein (BMP) signals in the Drosophila germline stem cells (GSCs) (Chen and McKearin, 2003). This model of niche function implies that the stem cell state is inherently unstable and must be actively maintained. However, in tissues varying from the Drosophila ovary and testis to the mouse intestine (Amoyel et al., 2016a; Haramis et al., 2004; He et al., 2004; Kirilly et al., 2011), disrupting certain signalling pathways results in the accumulation of stem cells, implying that stem cell differentiation requires active signalling.

In parallel to this model of stem cell fate in which niche or differentiation signals are instructive, analysis based on clonal lineage tracing has revealed that individual labelled stem cells can be lost from the niche and differentiate, while their neighbours fill the available space by symmetric division. Analysis of these clonal patterns shows that at the single cell level, self-renewal or differentiation decisions are stochastic (Klein and Simons, 2011). As well as this, lineage tracing of cells expressing markers of a more differentiated fate often results in stem cells being labelled, indicating that these cells can give rise to stem cells with long-term self-renewal potential. Such studies show that marker expression can fluctuate and does not reflect potential, giving rise to a more nuanced picture of what constitutes stem cell identity (Chatzeli and Simons, 2020; Clevers and Watt, 2018; Donati and Watt, 2015; Ge and Fuchs, 2018; Krieger and Simons, 2015; Tai et al., 2019). This work suggests that stem cell potential is distributed across a population of cells within which different states exist where some cells have a higher propensity, or bias, towards self-renewal and some towards differentiation. How stem cells transition between these states and out of the stem cell pool and the contribution of niche signals to these transitions is poorly understood.

To probe how and when stem cell daughters commit to differentiation, we study the somatic lineage in the Drosophila testis. The testis stem cell niche, called the hub, is composed of approximately 10-12 quiescent cells that support two stem cell populations (Fig. 1A) (Greenspan et al., 2015). GSCs adhere tightly to the hub and divide with invariantly oriented divisions, generating in most cases two daughters with asymmetric fates. The differentiating daughter, known as a gonialblast, undergoes 4 rounds of division with incomplete cytokinesis to form a cyst which matures into spermatocytes. The second population, somatic cyst stem cells (CySCs), give rise to post-mitotic differentiating cells, called cyst cells. These tightly associate with a gonialblast in a 2:1 ratio, ensheathing the developing cyst and eventually sealing off the germline from the environment (Fig. 1A) (Fairchild et al., 2015; Kiger et al., 2000; Shields et al., 2014; Tran et al., 2000). Coordination between the lineages and support from the soma are essential for germ cell differentiation and fertility (Kiger et al., 2000; Matunis et al., 1997; Smendziuk et al., 2015; Tran et al., 2000). Recent work has shown that this coordination is not regulated through synchronised stem cell divisions but instead at the level of co-differentiation of gonialblasts and CySC daughters, when encystment leads to abscission of the gonialblast from the GSC (Lenhart and DiNardo, 2015). How the two lineages temporally coordinate their differentiation is difficult to envisage if differentiation is simply a consequence of loss of niche signals. CySCs rely primarily on activity of the Janus Kinase and Signal Transducer and Activator of Transcription (JAK/STAT) pathway for self-renewal (Leatherman and Di Nardo, 2008), while additional signals including the Hedgehog (Hh), Hippo (Hpo), Mitogen-Activated Protein Kinase (MAPK) and Slit/Robo pathways regulate proliferation and niche occupancy (Amoyel et al., 2013, 2014, 2016b; Michel et al., 2012; Singh et al., 2016; Stine et al., 2014). BMPs produced by the hub and the CySCs maintain GSC self-renewal, and do so at least in part by directly repressing expression of the differentiation factor *bam,* supporting the model in which niche-derived signals repress differentiation (Kawase et al., 2004; Shivdasani and Ingham, 2003).

**Figure 1:**
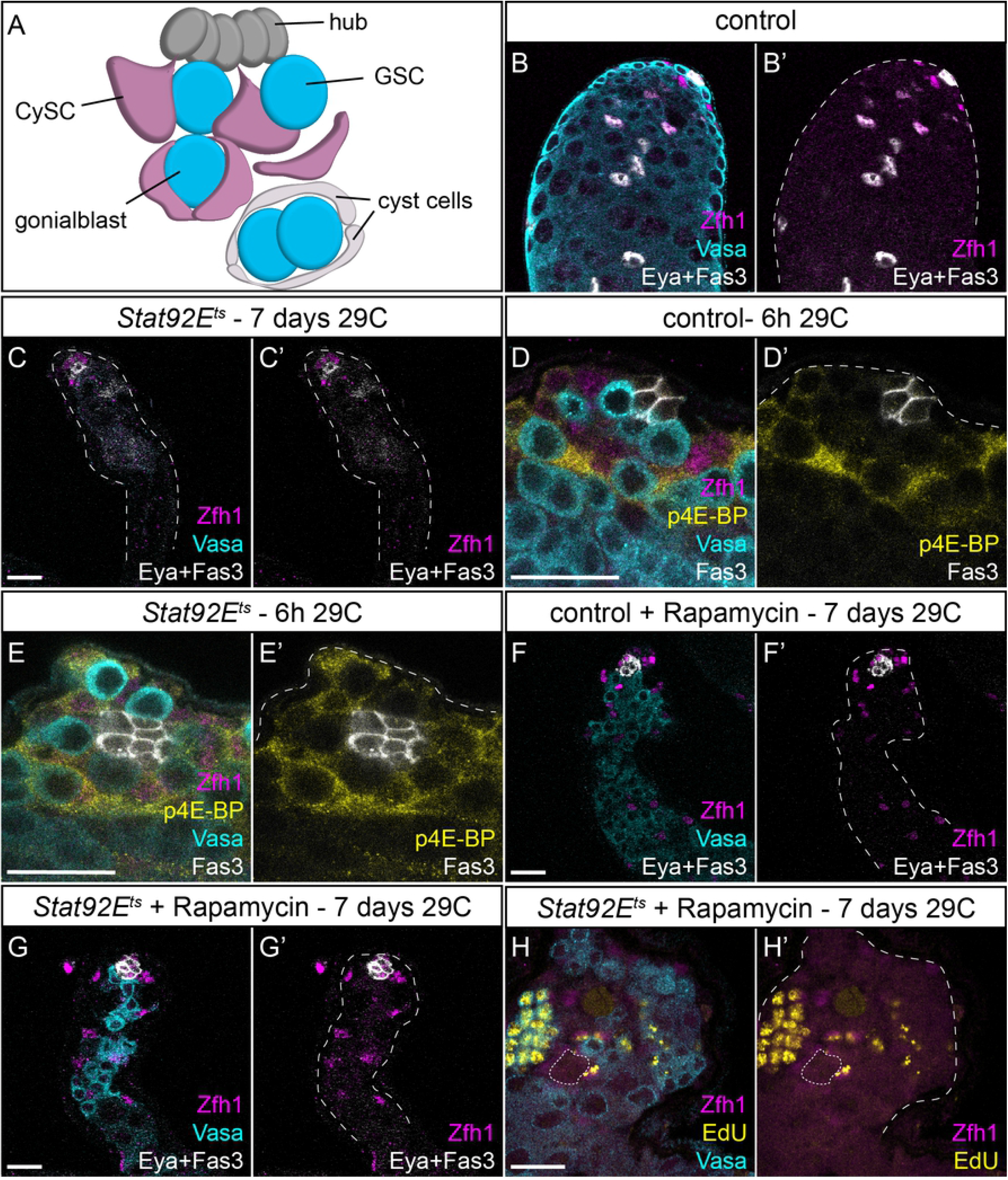
Tor is required for cyst cell differentiation in the absence of the self-renewal signal JAK/STAT. A. Diagram representing the apical tip of the Drosophila testis. Hub cells (dark grey) support two stem cell populations: GSCs (blue) and CySCs (magenta). GSCs divide to give rise to gonialblasts that are wrapped by two CySC daughters. These differentiate into cyst cells (white) and support the encysted germline. B,C. Control and *Stat92E^TS^* animals shifted to 29°C for 7 days showing loss of both CySCs and GSCs in the *Stat92E^TS^* mutant. Zfh1 (magenta) labels CySCs and early daughters, Vasa (cyan) labels the germline, and Eya and Fas3 (white) label differentiated cyst cells and the hub, respectively. D,E. p4E-BP is upregulated early in CySCs upon loss of JAK/STAT signalling. In controls (D), high levels of p4E-BP (yellow), are detected in Zfh1-positive (magenta) cells two rows away from the hub (Fas3-positive, white). In *Stat92E^TS^* mutants shifted for 6h, high p4E-BP is visible in CySCs contacting the hub. Vasa labels the germline in cyan and shows GSCs contacting the hub in both conditions. F-H. Rapamycin feeding inhibits CySC differentiation even in *Stat92E* mutants. In controls (F), rapamycin feeding results in Zfh1-expressing cells (magenta) being found throughout the testis, and no Eya-positive cells (white, compare to B). Rapamycin prevents the loss of Zfh1-expressing CySCs in *Stat92E^TS^* mutants after 7 days at 29°C (G), and no Eya-expressing cells are visible. H. EdU incorporation (yellow) is detected in Zfh1-expressing cells. Vasa (cyan) labels the germline, the hub is indicated with Fas3 expression or a dotted line. Scale bar in all panels represents 20 μm.

In the cyst lineage, however, clonal assays have revealed that CySC divisions produce equipotent daughters which stochastically self-renew or differentiate and that CySC replacement at the niche by neighbours is a frequent event (Amoyel et al., 2014). CySCs are identified by contact with the niche and progression through the cell cycle, while differentiated cells are post-mitotic; mitoses are only observed at the hub (Cheng et al., 2011; Hardy et al., 1979). CySCs express the transcription factor Zfh1, which is necessary and sufficient for self-renewal (Leatherman and Di Nardo, 2008). Zfh1 expression is downregulated during differentiation and cyst cells begin expressing the cyst cell marker Eyes Absent (Eya) (Fabrizio et al., 2003). Zfh1 expression labels a larger pool of cells than actually contact the hub, suggesting that CySC potential may extend beyond cells with direct hub contact. Consistently, although CySC divisions result in only one cell inheriting the attachment to the hub (Cheng et al., 2011), clonal experiments show that both daughters of a CySC division can gain access to the hub (Amoyel et al., 2014). Indeed, recent work tracing cells lacking hub cell contact showed that they can give rise to labelled CySCs (Chiang et al., 2017). Finally, cyst cell differentiation requires signalling through the Phosphoinositide 3-kinase (PI3K)/Target of Rapamycin (Tor) pathway (Amoyel et al., 2016a; Sênos Demarco et al., 2020; Tamirisa et al., 2018). Activity of PI3K and Tor is high in Zfh1-positive cells that do not contact the hub and CySCs deficient for Tor or PI3K pathway activity maintain Zfh1 expression and fail to express Eya, while clonal gain-of-function leads to differentiation (Amoyel et al., 2014, 2016a; Sênos Demarco et al., 2020). Thus, cyst cell differentiation is actively induced by signalling through PI3K/Tor, yet how the differentiation of cyst cells is temporally regulated to ensure coordination with the germline is not understood. Here we set out to understand how cyst cell differentiation is regulated by addressing the role of PI3K/Tor signalling in cyst cells. We show that CySC differentiation requires Tor activity even in the absence JAK/STAT signalling, implying that differentiation is not a default state in the absence of niche-derived self-renewal signals. Conversely, the sufficiency of PI3K/Tor to drive differentiation is relative: clonal increases in PI3K/Tor drive differentiation while lineage-wide increases do not, consistent with a model in which high PI3K/Tor activity primes cells for differentiation but is not sufficient for commitment. Furthermore, we show that the ability of increased PI3K/Tor to promote differentiation requires the presence of the germline. We propose that CySC differentiation into cyst cells occurs in reversible steps: loss of niche access and selfrenewal signals, followed by acquisition of differentiation competence driven by PI3K/Tor activity, before the final commitment to differentiation which occurs upon interaction with a gonialblast. Our work suggests that actively regulating differentiation in this way is an important mechanism for achieving coordination across lineages.

## Results

### Tor is required for cyst cell differentiation in the absence of JAK/STAT signalling

In the somatic cyst lineage of the testis, JAK/STAT signalling is both necessary and sufficient to maintain CySC fate, while PI3K/Tor activity is required for cyst cell differentiation (Amoyel et al., 2016a; Leatherman and Di Nardo, 2008). We sought to understand how differentiation is initiated upon withdrawal of JAK/STAT signalling, and whether Tor was required in this process.

We used a temperature-sensitive mutant for the sole Drosophila STAT, *Stat92E,* to deplete JAK/STAT activity in adult males. In testes from these mutants, as described previously (Brawley and Matunis, 2004; Leatherman and Dinardo, 2010), both GSCs and CySCs were rapidly lost to differentiation. By 7 days at the restrictive temperature (29°C), few, if any, Zfh1-positive CySCs were observed (Fig. 1C, compare to control in Fig. 1B). To characterise the events involved in initiating differentiation, we examined testes from *Stat92E^ts^* animals prior to the stem cells being lost to differentiation. Previous work has shown that 16h after shifting to the restrictive temperature, GSCs are still present (Leatherman and Dinardo, 2010). We used an antibody recognising phosphorylated eIF4E-Binding Protein (p4E-BP) to monitor Tor activity in *Stat92E^ts^* testes 6h after shifting to the restrictive temperature. At this stage, Zfh1-expressing CySCs and single round GSCs were still present (Fig. 1E), consistent with previous work showing that differentiation does not occur immediately upon loss of STAT activity. In control *Stat92E^ts^* heterozygous animals shifted to 29°C for 16h, p4E-BP was highly expressed in the second row of somatic cells from the hub, and lower in the first row of CySCs that directly contact the hub (Fig. 1D). In contrast, in *Slal92E”* mutant animals, the pattern of p4E-BP changed so that high p4E-BP was now detected in somatic cells directly adjacent to the hub (Fig. 1E). We counted 1.2 ± 0.2 (n=26) cells in contact with the hub with high p4E-BP in controls compared to 5.2 ± 0.4 (n=25) in *Stat92E^ts^* mutants (P<0.0001). As a control, in animals kept at the permissive temperature, there was no difference in the number of p4E-BP-positive cells at the hub between controls and mutants (2.0 ± 0.3 for *Stat92E^ts^* (n=11) compared to 2.2 ± 0.3 for *Stat92E^ts^* (n=6), P>0.9). These results show that Tor activity is induced in CySCs that will differentiate, and precedes detectable changes in morphology and marker expression.

Since previous work showed that Tor activity is required for differentiation (Amoyel et al., 2016a; Sênos Demarco et al., 2020) and since Tor activity is upregulated upon JAK/STAT loss, we asked whether Tor is required for the CySC differentiation that occurs following loss of Stat92E activity. In other words, we tested whether cyst cell differentiation is a default state in cells lacking JAK/STAT activity, or whether differentiation is induced by Tor. As shown above, loss of JAK/STAT by shifting *Stat92E^ts^* animals to the restrictive temperature for 7 days results in almost complete loss of CySCs and GSCs (Fig. 1B,C). Conversely, inhibiting Tor activity by feeding the inhibitor Rapamycin led to Zfh1-positive cells being found away from the hub which did not express Eya (Fig. 1F), indicating a block in differentiation. When *Stat92E^ts^* mutant animals were fed Rapamycin and shifted to the restrictive temperature, testes contained Zfh1-positive cells distant from the hub and few Eya-positive cells (Fig. 1G), indicating that these testes accumulated undifferentiated CySCs. In addition, Vasa-positive GSCs were present in these testes. To confirm that the ectopic Zfh1-positive cells are indeed CySCs, we assessed whether any cells incorporated the nucleotide analogue EdU, as CySCs are the only proliferating somatic cells in the testis. We observed EdU incorporation in Zfh1-positive cells in *Stat92E^ts^* animals fed with Rapamycin (Fig. 1H), indicating that Rapamycin feeding maintained CySCs even in the absence of JAK/STAT activity.

Altogether, these results demonstrate that cyst cell differentiation requires active signalling through Tor, even in the absence of the main self-renewal factor, JAK/STAT. This finding suggests that CySC identity can be maintained in the absence of niche signals, as long as differentiation signals are also inhibited.

### Clonal increases in Tor activity promote cyst cell differentiation

These results together with previously published work (Amoyel et al., 2016a; Sênos Demarco et al., 2020; Tamirisa et al., 2018) show that Tor activity is absolutely necessary for cyst cell differentiation. We next asked whether increases in Tor activity directly induce differentiation or whether it is merely permissive for differentiation. To this end, we used mitotic recombination to generate single cell CySC clones with gain-of-function mutations in the PI3K/Tor pathway.

Since CySCs are the only mitotic somatic cells, all labelled somatic cells are the result of a CySC division. We examined clones at 2 days post clone induction (dpci) to verify induction rates, and at 7 dpci to assess the persistence of these clones, which directly reflects the self-renewal and differentiation capacity of the labelled cells.

In controls, clones were reliably induced at 2 dpci (Fig. 2A,E), and many of these clones were present a week after induction (Fig. 2B,F). Clones at 7dpci consisted of both CySCs, identified as the first row of Traffic Jam (Tj)-positive somatic cells from the hub (Fig. 2B,F arrowheads), and differentiated cyst cells, which were displaced from the hub and grew larger (Fig. 2B,F arrows). Clones mutant for genes encoding components of the Tor repressive Tuberous Sclerosis Complex (TSC), *Tsc1* or *gigas (gig,* the Drosophila homologue of *Tsc2)* were also recovered at 2 dpci (Fig. 2C,G, arrowheads), indicating that mutant CySCs could be induced. However, by 7 dpci, labelled CySC clones were rarely recovered (Fig. 2D,H,I, P<0.0001 for *Tsc1* compared to control, P<0.005 for *gig* compared to control), implying that the mutant cells had differentiated. Indeed, some testes with no CySC clones contained labelled cells distant from the hub, consistent with a mutant CySC that had differentiated and left the niche (Fig. 2D,H, arrows). We obtained similar results when generating clones mutant for the PI3K inhibitor, *Pten* (see below, Fig. 4C,D,I), confirming previous observations that *Pten,* and *Tsc1* mutant and wild type clones overexpressing the catalytic subunit of PI3K, Dp110, were not maintained at the niche and differentiated (Amoyel et al., 2014).

**Figure 2:**
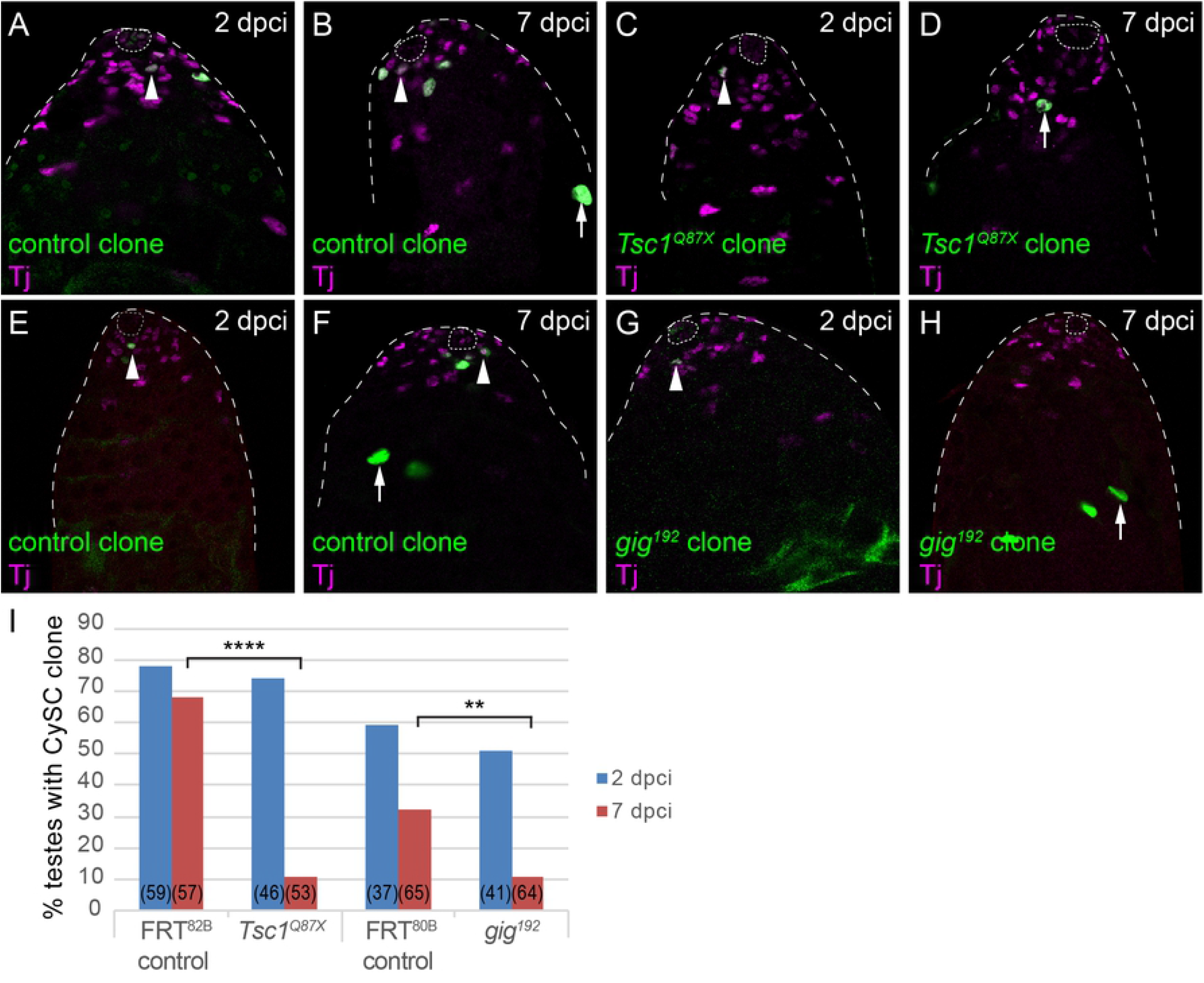
Clonal gain-of-function in Tor activity is sufficient to induce CySCs to differentiate. A-H. Positively-labelled CySC clones generated by MARCM. Control clones generated with *FRT^82B^* (A,B) or *FRT^80B^* (E,F) were observed at 2 days post clone induction (dpci) and at 7 dpci to assess clone induction and maintenance rates, respectively. CySCs (arrowheads) were identified as cells expressing Tj (magenta) and in the first row from the hub, differentiated cyst cells (arrows) have larger nuclei and are found away from the hub. *Tsc1* mutant (C,D) or *gig* (Tsc2, G,H) CySC clones could be induced and are present at 2 dpci (C,G) but were rarely observed by 7 dpci (D,H), where differentiated cells were labelled. The hub is outlined with a dotted line. Scale bar in all panels represents 20 μm. I. Graph showing the percentage of testes containing a CySC clone at 2 (blue) and 7 (red) dpci. The number of testes examined is showed in parentheses for each column. Statistical significance was determined using Fisher’s exact test, **** denotes P<0.0001, ** denotes P<0.005.

In sum, CySC clones with hyper-activated PI3K or Tor signalling self-renew less well than control CySC clones, and are lost from the niche through differentiation. This implies that PI3K/Tor activity is either deleterious for self-renewal, or is sufficient to promote differentiation. Since the results described in prior work (Amoyel et al., 2016a; Sênos Demarco et al., 2020; Tamirisa et al., 2018) and in Figure 1 argue that Tor is necessary for differentiation, the most parsimonious interpretation is that clonally increasing PI3K/Tor activity induces CySCs to differentiate into cyst cells.

### Lineage-wide increases in Tor activity do not affect CySCs

The data presented so far appeared to support a model in which, as daughters of a CySC division leave the hub and are out of the range of JAK/STAT signalling, they upregulate PI3K/Tor activity to differentiate. This is consistent with the observation that high levels of the Tor target p4E-BP are detected in the second row of Zfh1-expressing cells (Fig. 1D and (Amoyel et al., 2016a; Sênos Demarco et al., 2020)). However, although CySCs divide such that only one daughter cell inherits the hub attachment (Cheng et al., 2011), clonal analysis showed that both daughter cells are equipotent (Amoyel et al., 2014), suggesting that cells away from the hub maintain the ability to self-renew. Moreover, lineage tracing using *spict-Gal4,* which is expressed in Zfh1-positive cells in the second row, indicate that these cells do give rise to CySCs (Chiang et al., 2017), raising the possibility that cells with high PI3K/Tor do not necessarily differentiate. We confirmed that indeed, *spict-Gal4-driven* GFP and high levels of p4E-BP do indeed colocalise in Zfh1-expressing cells (Fig. S1, arrows). Thus, cells with high PI3K/Tor are not necessarily committed to differentiate.

As a further test for the sufficiency of increased PI3K/Tor in promoting CySC differentiation, we hyper-activated signalling in all CySCs using the *traffic jam (tj)-Gal4* driver. We predicted that, if increasing signalling drives differentiation, many or most CySCs would be lost to differentiation, resulting in testes containing few or no Zfh1-expressing cells, similar to loss-of-function of JAK/STAT signalling. Surprisingly, over-expression of Dp110 or knockdown of *Tsc1* using *tj-Gal4* resulted in testes that were indistinguishable from control (compare Fig. 3A,B and C). There was no significant change in the number of Zfh1-positive and Eya-negative CySCs (45±1.1 in control compared to 41.9±1.5 and 45.5±2.4 in Dp110-expressing or *Tsc1* knockdown testes, respectively, n=19,18,11, P<0.25). To rule out experimental artefacts, we tested whether these constructs could induce ectopic PI3K and Tor activity. First, we used *engrailed-Gal4* to drive expression of *UAS-Dp110* or *UAS-Tsc1 RNAi* in the posterior compartment of larval wing imaginal discs, where PI3K and Tor are well-established regulators of growth during development. Compared to the anterior compartment of the same discs, Dp110 over-expression and *Tsc1* knockdown both led to reduced density of cell nuclei, reflecting increased growth (Fig. S2A-D). We also used established readouts for the activity of both pathways: phosphorylated Akt1 (pAkt) for PI3K signalling, and phosphorylated S6 Kinase (pS6k) for Tor signalling. Overexpressing Dp110 in wing discs led to a robust increase in pAkt (Fig S2B) and knockdown of *Tsc1* resulted in increased pS6k staining in the posterior compartment (Fig. S2D). Finally, we tested for increased pathway activity in testes under these conditions. *Tsc1* knock down or Dp110 over-expression in the cyst lineage resulted in increased p4E-BP staining (Fig 3E,F, compare with 3D), indicating that these manipulations elevate Tor activity, even if they do not affect the number of CySCs.

**Figure 3:**
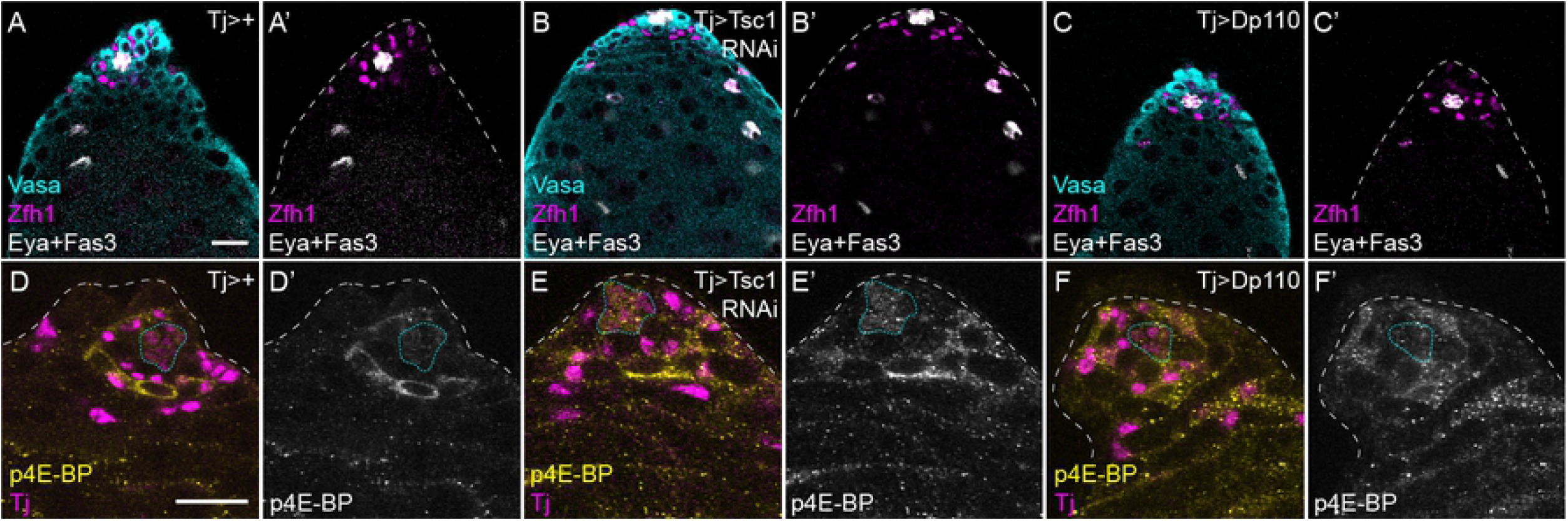
Lineage-wide PI3K/Tor hyperactivation does not induce CySC loss. Control (A,D), Tsc1 knockdown (B,E) or Dp110 over-expression (C,F) in the somatic lineage using *tj-Gal4.* A-C. Zfh1 (magenta) labels CySCs and early daughters, Eya (white) labels differentiated cyst cells and Fas3 (white) labels the hub. Germ cells are labelled with Vasa in cyan. No differences in the numbers Zfh1-expressing CySCs are observed upon PI3K or Tor hyperactivation. D-F. p4E-BP staining (yellow) is increased upon Tsc1 knockdown (E) or Dp110 over-expression (F). Tj labels the somatic lineage (magenta). The hub is outlined with a blue dotted line. Scale bar in all panels represents 20 μm.

We therefore conclude that increasing PI3K or Tor activity in CySCs is sufficient to drive differentiation when such manipulations are carried out in clones, but not in the entire CySC population.

### Differentiation of CySCs with increased PI3K activity depends on signalling in neighbouring cells

The experiments described above suggest a model in which PI3K/Tor activity induces a permissive state for differentiation, but does not commit cells to differentiation. We propose that PI3K/Tor activity acts as a licensing factor to permit differentiation, and therefore increases the likelihood of differentiation, but upon exposure to hub-derived signals, cells experiencing high PI3K/Tor can self-renew. This model predicts that in clonal experiments, increasing PI3K/Tor results in an increased probability of differentiation and eventual loss of the clone from the stem cell pool, while in lineage-wide gain-of-function, all cells are licensed to differentiate but only a proportion will commit to a differentiated fate.

We took two approaches to test the model that PI3K/Tor signalling primes cells for differentiation. First, one prediction is that the differentiation of *Pten* mutant clones is due to higher PI3K/Tor signal in the mutant clones causing them to remain in a permissive state for differentiation, and not an intrinsic inability to self-renew. We reasoned that the maintenance of *Pten* mutant CySC clones could be rescued by increasing PI3K levels in all CySCs, thereby equalising the differentiation likelihood between *Pten* mutants and neighbours. We generated negatively-marked clones, either wild type or mutant for *Pten*, and assessed whether these clones were observed at 2 dpci and 7 dpci. As previously, control Tj-positive clones lacking GFP were observed in the first row from the hub at both 2 and 7 dpci, indicating that CySC clones were induced and maintained over the course of the experiment (Fig. 4A,B). In contrast, and consistent with published data (Amoyel et al., 2014), *Pten* mutant clones, although induced robustly (Fig. 4C), were rarely recovered by 7 dpci (Fig. 4D,I, P<0.0001). When we used the C587-Gal4 driver to over-express Dp110 in CySCs, wild type clones were recovered at similar rates to control (Fig. 4E,F,I, P<0.7). However, *Pten* mutant clones induced in testes where Dp110 was expressed with *tj-Gal4* were recovered as frequently as wild type clones at 7 dpci (Fig. 4G-I, P<0.06) and were significantly rescued compared to *Pten* mutant clones in a control background (P<0.0001). These data indicate that *Pten* mutant CySCs maintain the ability to self-renew but that they are normally prevented from doing so by an increased tendency to differentiate compared to neighbouring cells.

**Figure 4:**
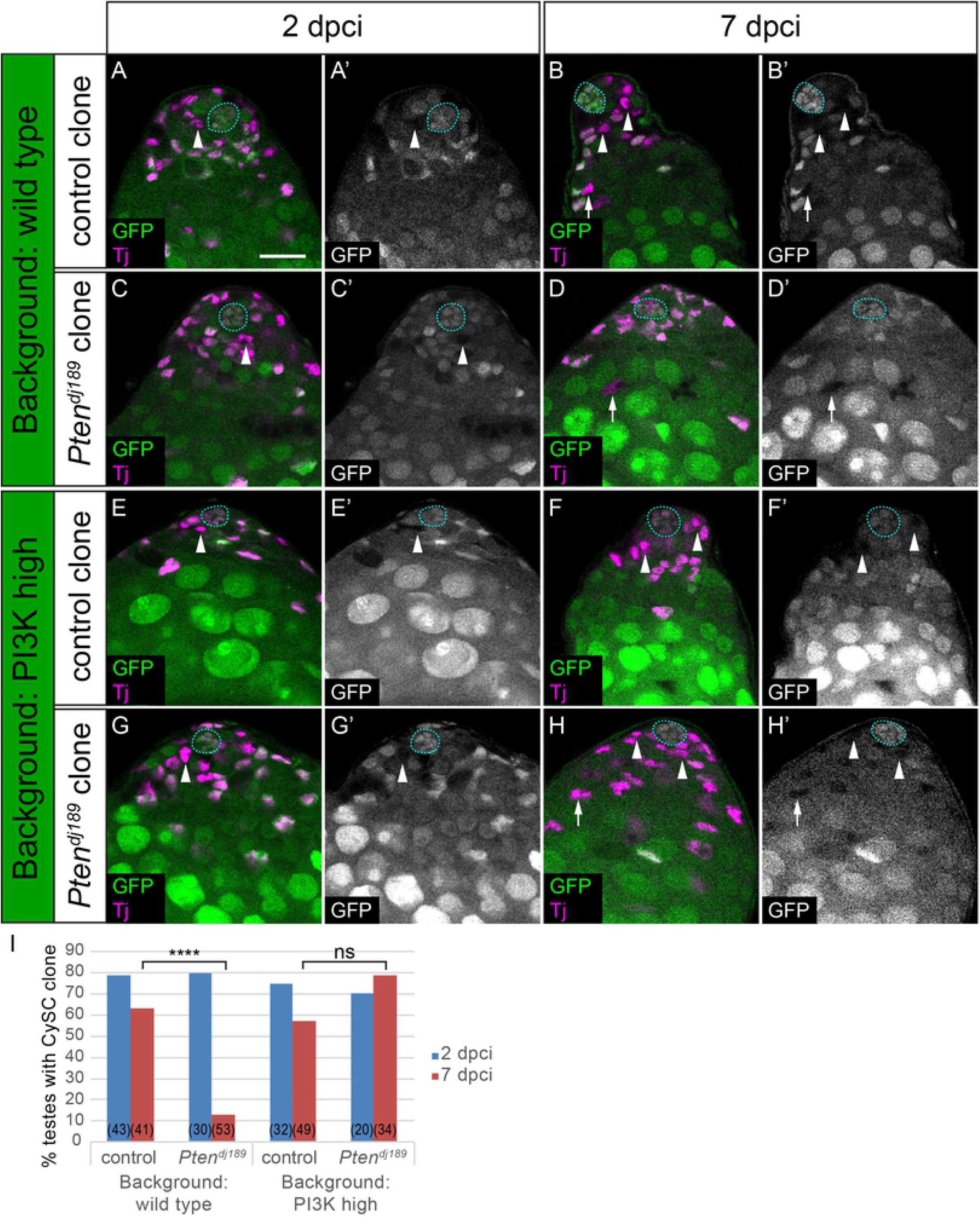
Differentiation of PI3K-high CySCs depends on signalling levels in neighbouring cells. Control (A,B,E,F) and *Pten* mutant (C,D,G,H) clones induced by mitotic recombination and marked by loss of GFP expression, generated either in a control background *(C587-Gal4>+,* AD) or in a background with elevated PI3K activity *(C587-Gal4>Dp110,* E-H). CySCs (indicated with arrowheads) were identified as Tj-positive (magenta) nuclei one cell diameter from the hub (outlined with a blue dotted line). While *Pten* mutant CySC clones were rarely recovered at 7 dpci, mutant cyst cells were observed (D, arrow) indicating differentiation of the clones. By contrast, *Pten* mutants self-renewed and CySC clones were recovered when PI3K activity was elevated (H, arrowheads). Scale bar in all panels represents 20 μm. I. Graph showing the percentage of testes containing a CySC clone at 2 (blue) and 7 (red) dpci. The number of testes examined is showed in parentheses for each column. Statistical significance was determined using Fisher’s exact test, **** denotes P<0.0001, ns: not significant.

As a second test of the hypothesis that PI3K/Tor activity acts to licence cells for differentiation, we generated mosaic testes containing cells with defined levels of PI3K activity. To ensure that both cell types can both self-renew and differentiate, we generated clones that were wild type in a background where all cells had elevated PI3K activity (Fig. 5A, and see methods for how clones were generated). In this situation, as shown in Fig. 3, Dp110 over-expressing CySCs had no detectable phenotype and self-renewed well. However, we predicted that if PI3K levels determined the ability of cells to differentiate, wild type clones would be less likely to differentiate than Dp110-expressing cells, and that over time, wild type cells should come to dominate the CySC pool. We expressed Dp110 together with GFP under the control of *tj-Gal4* and induced clones in this background that expressed the Gal4 inhibitor, Gal80. These clones had no Gal4 activity and were therefore wild type for PI3K, and were labelled by loss of GFP expression. When we generated Gal80-expressing wild type clones in testes expressing only GFP under *tj-Gal4* control, we observed a mix of GFP-positive and negative cells with a broad distribution of clone sizes (Fig. 5B,D), consistent with previous observations that CySCs compete stochastically with each other for niche occupancy, with each stem cell having an equal chance of outcompeting its neighbour or of being displaced from the niche (Amoyel et al., 2014). When we induced Gal80-expressing clones in testes expressing both GFP and Dp110 under *tj-Gal4* control, many more GFP-negative cells were observed at 14 dpci (Fig. 5C,D, P<0.0006). This was not due to increased clone induction rates, as similar numbers of GFP-negative cells were present at 2 dpci (14.7 ± 1.6 GFP-negative cells in control compared to 13.2 ± 2.8 in a Dp110-expressing background, n=29 and 13, respectively, P<0.58). Thus, the ability of Gal80 clones with wild type levels of PI3K to self-renew was significantly affected by PI3K levels in neighbouring cells: in control conditions, these clones have no obvious advantage; however, when surrounded by PI3K-high cells, they are more likely to self-renew and colonise the niche.

**Figure 5:**
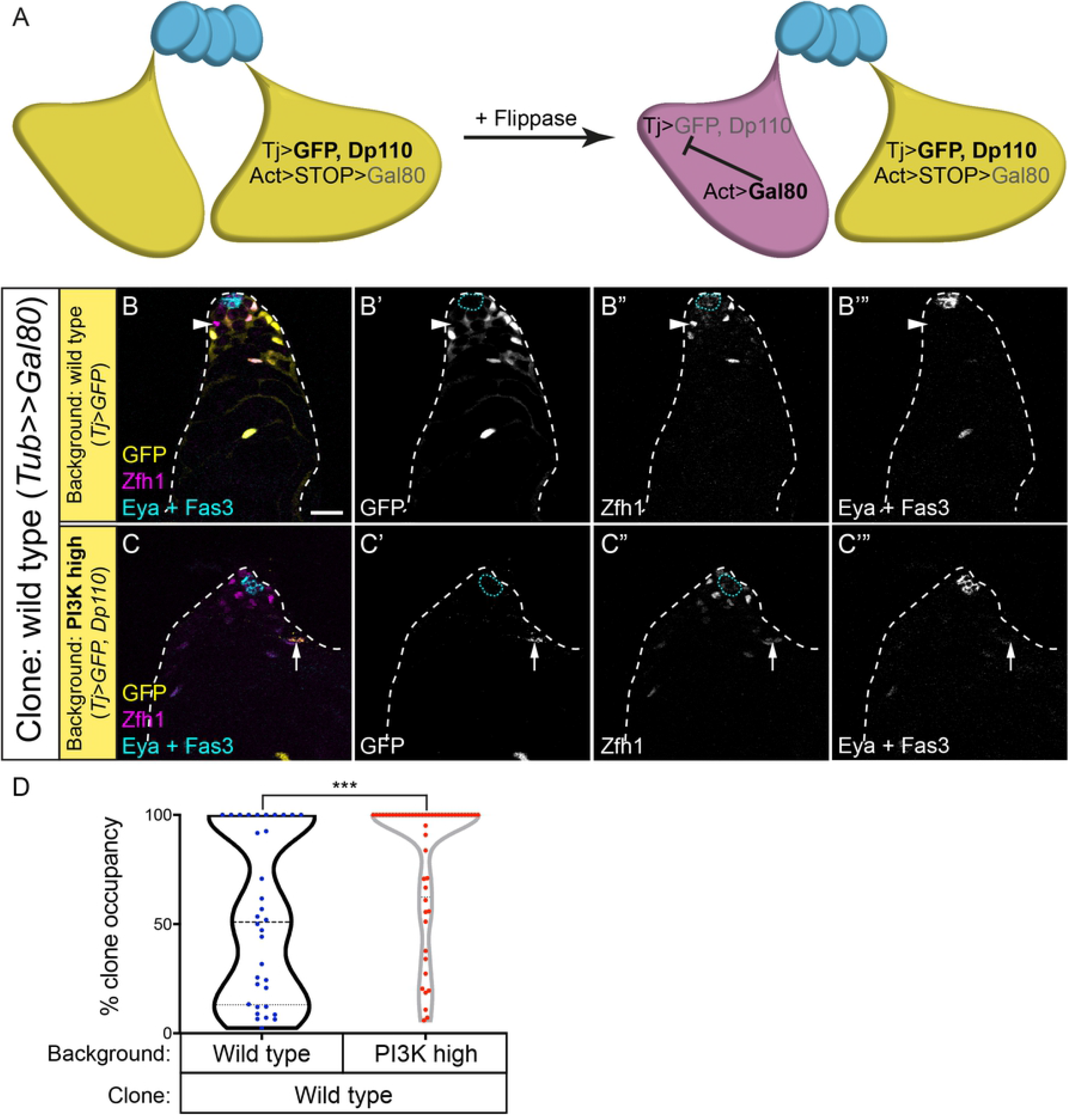
Differences in neighbour PI3K activity levels influences the niche occupancy of wild type clones. A. Schematic explaining the genotypes of clones and background. All CySCs express GFP (yellow) and Dp110 under *tj-Gal4* control while a flipout cassette with an Actin promoter driving Gal80 expression is inactive due to a STOP sequence. Flippase induction excises the STOP cassette, resulting in Gal80 expression which inhibits the activity of Gal4. The resulting cell (magenta) is GFP-negative and does not over-express Dp110. B,C. Wild type Gal80-expressing clones were generated in the cyst lineage that lacked GFP (yellow, single channels in B’,C’). CySCs were labelled with Zfh1 (magenta, single channels in B”,C”), and the hub and cyst cells with Fas3 and Eya respectively (white, single channels in B”’,C”’). B. At 14 dpci, wild type CySC clones (arrowhead) were observed surrounded by GFP-expressing CySCs. C. When GFP-expressing cells also had high PI3K activity, wild type CySC clones occupied a greater proportion of the stem cell pool, and often replaced all GFP-expressing CySCs. Arrow in C points to a GFP-expressing, Eya-positive differentiating cyst cell. Scale bar represents 20 μm. D. Quantification of the clone occupancy for wild type clones in different backgrounds. Clone occupancy is the number of GFP-negative cells as a proportion of all Zfh1-positive cells. Statistical significance was determined using a Mann-Whitney test, *** denotes P<0.0005.

### Germ cells promote cyst lineage differentiation

The experiments described above indicate that PI3K/Tor signalling is required to allow CySCs to differentiate, but that it is not sufficient to commit cells to differentiation. This implies that differentiation itself requires an additional input, and that only a subset of the cells that are primed receive this and go on to differentiate. In the testis, cyst cells co-differentiate with GSC daughters, gonialblasts, such that two cyst cells encapsulate one goniablast. Gonialblasts, therefore, are a good candidate to be the limiting resource for cyst cell differentiation.

First, we tested whether the presence of a germline affected cyst lineage differentiation. Previous work has shown that agametic testes display ectopic proliferating somatic cells, consistent with excess undifferentiated CySCs being present (Gonczy and DiNardo, 1996). We generated testes in which the germline was forced to undergo differentiation, by expressing the germ cell differentiation factor Bam under control of the GSC and early germ cell driver *nanos (nos)-Gal4* (Sheng et al., 2009). Bam expression in the germline resulted in variably penetrant germ cell loss from the niche. In testes completely lacking a germline, we observed ectopic Zfh1-expressing cells many cell diameters away from the hub, and few Eya-positive cells (Fig. 6A,B). Moreover, we detected many Zfh1-positive cells that were proliferating, as assessed by EdU incorporation (Fig. 6B). These data show that loss of the germline results in ectopic cells with CySC-like fate, consistent with published work implicating germ cells in promoting cyst cell differentiation (Gonczy and DiNardo, 1996).

**Figure 6:**
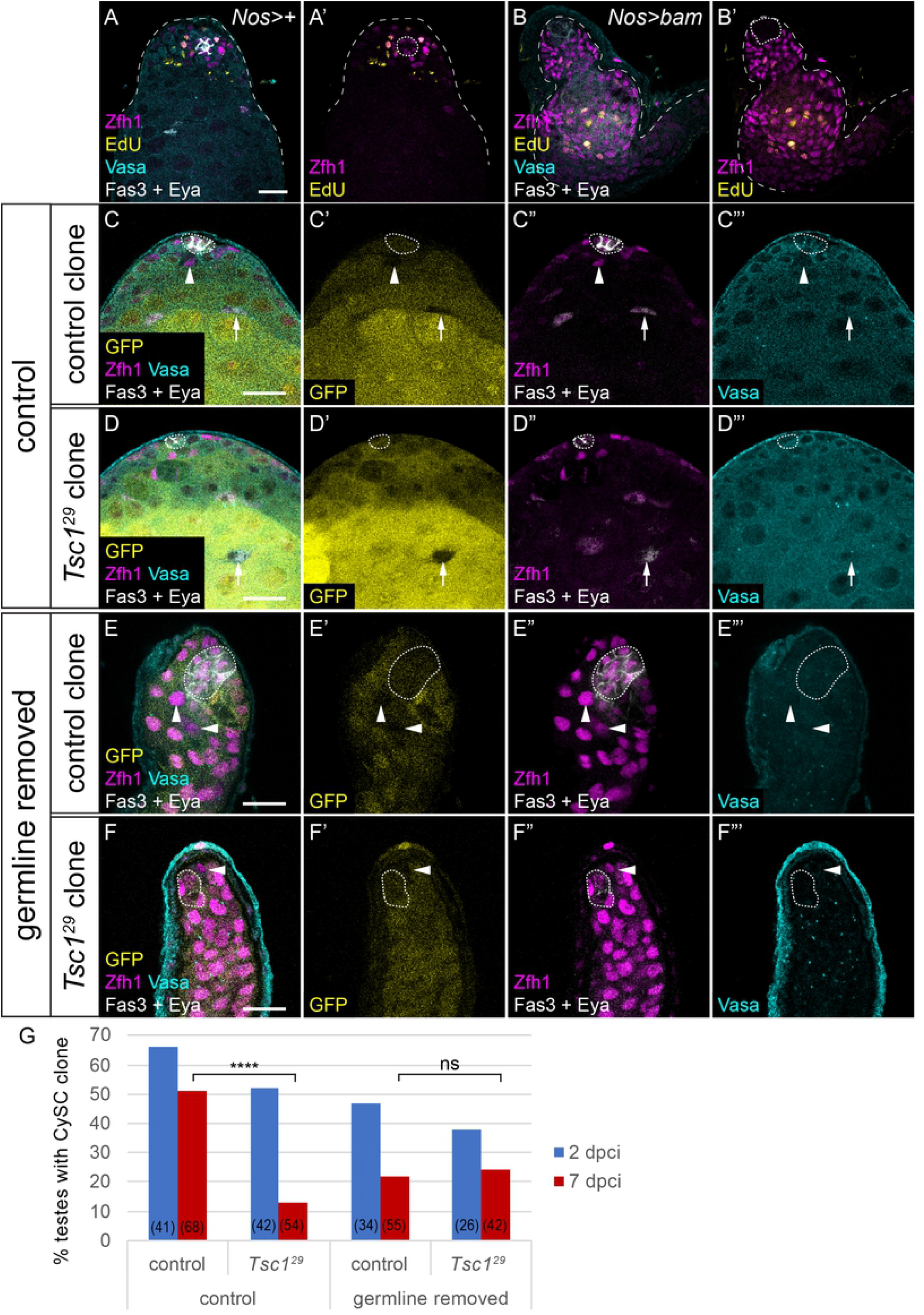
The germline is required for differential PI3K activity to induce differentiation. A,B. Control (A) and germline-ablated (B) testes, showing Zfh1 (magenta) to label CySCs and early daughters, Fas3 and Eya (white) to label the hub and differentiated cyst cells respectively, Vasa (cyan) to label the germline and EdU (yellow) labelling cells replicating DNA. Germline ablation results in loss of Vasa staining, expansion of Zfh1 away from the hub and EdU incorporation in somatic cells away from the hub. C-F. Control (C,E) or *Tsc1* mutant (D,F) negatively-marked clones at 7 dpci. Clones are identified by the absence of GFP (yellow, single channel C’-F’). In testes where the germline (Vasa-positive, cyan, single channel C’”-F”’) is present control CySC clones (C) are recovered and contain both Zfh1-expressing CySCs (arrowheads, magenta) and Eya-positive cyst cells (arrows, white), while *Tsc1* mutant CySCs (D) are rarely observed. By contrast, when the germline is absent (E,F), control and *Tsc1* mutant clones are recovered at similar rates, and both consist of only Zfh1-expressing cells. Scale bar in all panels represents 20 μm. G. Graph showing the percentage of testes containing a CySC clone at 2 (blue) and 7 (red) dpci. The number of testes examined is showed in parentheses for each column. Statistical significance was determined using Fisher’s exact test, **** denotes P<0.0001, ns: not significant.

To test the hypothesis that CySC daughter cells are competing for gonialblasts to co-differentiate with, we induced clones with a gain-of-function mutation in the PI3K/Tor pathway in agametic testes. If the role of PI3K/Tor is indeed to prime CySC daughter cells to encapsulate a gonialblast, the lack of gonialblasts would result in clones with elevated PI3K/Tor remaining undifferentiated. Conversely, if elevating PI3K/Tor was sufficient to drive differentiation independently of the germline, these clones should differentiate, similar to PI3K/Tor gain-of-function clones in wild type testes (Fig. 2). Clones mutant for the Tor pathway inhibitor *Tsc1* were readily induced and observed at 2 dpci (Fig. 6D,G), but differentiated rapidly and few CySC-containing clones were recovered at 7 dpci, compared to control clones (Fig. 6C,G, P<0.0001). However, when we generated *Tsc1* mutant clones in testes where the germline underwent forced differentiation, these clones were now recovered at similar rates to control clones (Fig. 6E-G, P<0.8). Thus, in the absence of germ cells to encyst, CySCs with elevated Tor activity can self-renew similarly to control CySCs. These data imply that Tor does not autonomously promote differentiation, but, together with our previous experiments, show that Tor activity regulates the ability of individual CySC daughter cells to co-differentiate with the germline.

## Discussion

The data presented here suggest that cyst cell differentiation is an induced state, and that CySC daughter cells need multiple inputs in order to commit to differentiation (Fig. 7). We propose that upon losing access to niche-derived JAK/STAT pathway ligands, CySCs can be primed to differentiate by high levels of PI3K/Tor activity, but do not commit to differentiation until they associate with a gonialbast. Primed CySCs may return to the niche and continue to self-renew if they do not receive this secondary signal. This mechanism to limit differentiation of cyst cells ensures the temporal coordination of cyst cell and gonialblast differentiation.

**Figure 7:**
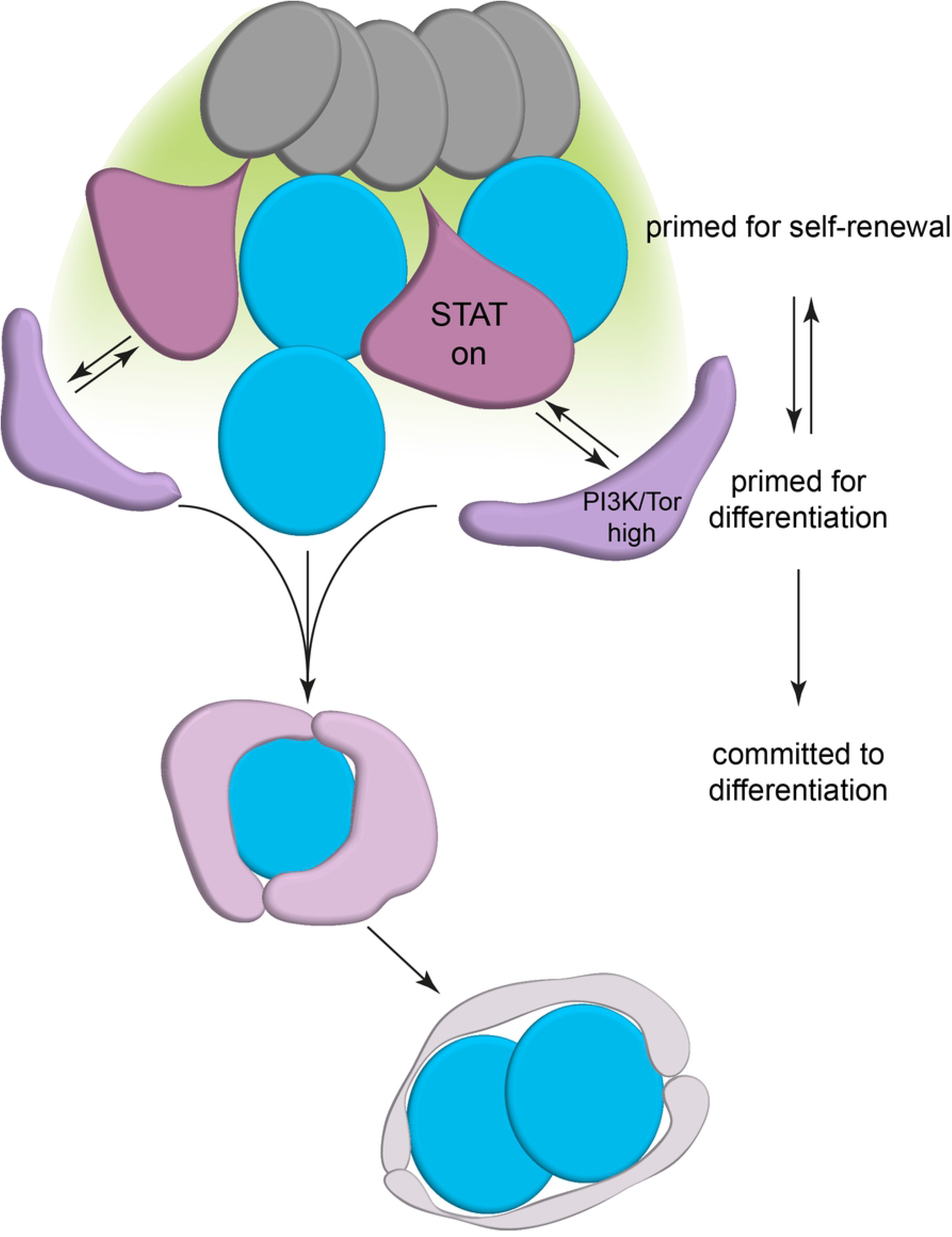
Model of CySC differentiation. CySCs (purple) at the hub (dark grey) receive self-renewal signals (indicated by a green gradient) and are primed for self-renewal. Upon losing access to the hub and increased PI3K/Tor activity, CySC daughter cells become primed for differentiation, and can interact with goniablasts that arise from GSC divisions. Differentiation priming is reversible, and cells with high PI3K/Tor maintain stem cell potential. Upon interaction with a gonialbast, a cyst is formed and the cells co-differentiate.

While much emphasis has been placed on the role of the niche in maintaining stem cell selfrenewal, recent work indicates that in the ovary, a separate niche promotes germ cell progression through differentiation (Kirilly et al., 2011; Wang et al., 2015). Similarly, in the vertebrate intestine, BMP inhibition results in ectopic stem-like cells, indicating a role for BMP signalling in promoting differentiation (Haramis et al., 2004; He et al., 2004). In both these instances, however, loss of the differentiation signal results in increased self-renewal signalling. Indeed, in the fly ovary, reducing the genetic dose of the GSC self-renewal ligand *dpp* could suppress the accumulation of undifferentiated germ cells caused by disrupting the differentiation niche (Wang et al., 2015). By contrast, we show here that Tor is required independently of JAK/STAT signalling to promote cyst cell differentiation, confirming previous work which had shown no upregulation of STAT activity in ectopic CySC-like cells observed upon PI3K/Tor loss-of-function (Amoyel et al., 2016a). CySCs deprived of both self-renewal and differentiation-promoting signals maintain a stable stem-like fate, implying that differentiation is not an inevitable consequence of losing access to niche-derived signals. This situation is reminiscent of that described for embryonic stem cells (ESCs), in which self-renewal signals are not necessary for stem cell maintenance if differentiation signals are inhibited (Ying et al., 2008). Thus, the ground state of ESCs is to self-renew in the absence of extrinsic signals, and we propose that CySCs similarly have a ground state favourable to self-renewal. This finding suggests that fate decisions may be regulated similarly in embryonic and adult stem cells.

Our results indicate that PI3K/Tor is not simply sufficient for differentiation, however, raising the question as to its role. One attractive model is that PI3K/Tor activity “primes” CySC daughter cells to differentiate, such that Zfh1-positive cells with high PI3K/Tor are able to exit the stem cell pool, but require other signals to do so. We show that cells experiencing high PI3K/Tor activity can go on to maintain CySC fate and self-renew, consistent with the observation that Zfh1-expressing cells in the second row from the hub can give rise to CySCs over time (Chiang et al., 2017). These cells, although more likely to differentiate, are not necessarily committed to differentiation, and can return to positions adjacent to the hub where PI3K/Tor signalling is lower. This model implies that CySCs exist in multiple states, where some are more biased towards self-renewal, and others towards differentiation, but that cells can transit reversibly between these states (Chatzeli and Simons, 2020; Enver et al., 2009). As such, the effective stem cell pool consists of both cells directly contacting the hub and those behind them, and the role of PI3K/Tor signalling is to promote the transition to a state biased towards differentiation. How it does so, and what its relevant targets are, is still unclear, although recent work indicates that suppression of autophagy by Tor is important for cyst cell differentiation (Sênos Demarco et al., 2020). Intriguingly, PI3K activity inhibits clonal expansion in mammalian epidermis by promoting progenitor differentiation, but promotes ectopic proliferation and hyperplasia when activated in a widespread manner, suggesting that similar mechanisms may be at play in control of epidermal progenitor differentiation (Murayama et al., 2007; Ying et al., 2018).

Finally, we present evidence that exit from the cyst stem cell pool depends on the presence of the germline and that PI3K/Tor activity does not regulate this step in differentiation. Previous work had shown that the germline is required to promote cell cycle exit and fate progression in the cyst lineage (Gonczy and DiNardo, 1996), and conversely, that the encystment of each gonialblast by two cyst cells is required for germ cell differentiation (Kiger et al., 2000; Tran et al., 2000). How this coordination and ratio are achieved is still poorly understood; recent work showed that the two stem cells do not synchronise their divisions to produce daughters to co-differentiate (Lenhart and DiNardo, 2015). Rather, the abscission, and subsequent differentiation, of the gonialblast from its sibling GSC depends on encystment by cyst cells. The data presented here show that gonialblasts are required for cyst cell differentiation even when Tor is elevated, suggesting that in normal conditions cyst cell differentiation is delayed until a cyst composed of two cyst cells and one gonialblast is assembled. This is consistent with the observation that differentiated cyst cells are always found associated with germ cell cysts. Thus, regulated differentiation of both lineages is essential in order to achieve the coordinated assembly of a cyst. We speculate that PI3K/Tor may prime CySC daughters to interact with neighbouring gonialblasts and promote assembly of this 3-cell cyst. This would explain why relatively high PI3K/Tor activity levels induce CySCs to differentiate, by promoting cyst cell-gonialblast interactions. In the absence of a neighbouring gonialblast, a cell with high PI3K/Tor may come under the influence of niche signals and continue to self-renew. What the final signal promoting the irreversible differentiation of the cyst and exit of the CySC daughters from the stem cell pool is currently unknown. Our model is remarkably similar to that proposed in the case of the mammalian testis, where the germline stem cell pool consists of two populations, one of which is biased towards self-renewal and another towards differentiation, but where individual cells can reversibly move between the populations, until they commit to differentiation (Hara et al., 2014; Nakagawa et al., 2010). This model provides an elegant solution to ensuring that the balance of self-renewal and differentiation is maintained when the differentiation cue is external to the lineage, in the case of the testis, induced by periodic signals of the seminiferous cycle (Yoshida, 2018). Our data suggest that, in addition, segregating the signals controlling licensing and commitment to differentiation may ensure the robust coordination of multiple lineages for proper tissue function.

## Materials and methods

### Fly stocks and husbandry

We used the following fly stocks: Oregon-R; *Stat92E^F^; Stat92E^85C9^; FRT^82B^, Tsc1^Q87X^; FRT^82B^, Tsc1^29^; gig^192^, FRT^80B^; Pten^dj189^, FRT^40A^; UAS-Tsc1 RNAi* (BDSC#54034); *UAS-Dp110* (gift of L. Johnston); *UAS-bam::GFP* (gift of R. Lehmann); *tj-Gal4* (Kyoto DGRC#104055); *nos-Gal4-VP16* (gift of R. Lehmann); *C587-Gal4* (gift of R. Lehmann); *spict-Gal4* (Kyoto DGRC#112900); *en-Gal4* (gift of L. Johnston); *tub>stop>Gal80* (BDSC#39213).

Rapamycin feeding was achieved by keeping flies on fly food to which 100 μl of a 4 mM rapamycin stock solution in ethanol was added and air dried. Flies were transferred to fresh vials every 2 days.

Flies were raised at 25°C, except *tj-Gal4* and *C587-Gal4* crosses which were raised at 18°C and adult flies of the correct genotype were maintained at 29°C for 10 days to achieve maximum Gal4 activity.

Positively-marked clones were induced using the MARCM technique (Lee and Luo, 1999) with a single 1h-long heat shock at 37°C. Gal80 clones were generated using a flipout cassette in which GAL80 is expressed upon recombination-mediated excision of a stop sequence (Bohm et al., 2010). Clones were induced using hs-FLP^12^ (BDSC#1929) and with a 30 minute-heat shock at 37°C to maintain low induction rates.

For a full list of genotypes by figure, see Table S1.

### Immunohistochemistry

Following dissection, testes were fixed in 4% paraformaldehyde in PBS for 15 minutes, then permeabilised twice for 30 minutes in PBS with 0.5% Triton X-100. Samples were the blocked in PBS, 0.2% Triton X-100, 1% Bovine Serum Albumin (PBTB), before being incubated overnight with primary antibodies at 4°C, washed twice for 30 minutes in PBTB, and incubated at room temperature for 2 hours with secondary antibodies. After two washes in PBS, 0.2% Triton X-100 for 30 minutes, testes were mounted on slides with Vectashield medium (Vector labs), and imaged using a Zeiss LSM880, or a Leica Sp8 confocal microscope. To detect phosphorylated epitopes, we dissected and fixed in buffer containing phosphatase inhibitors as previously described (Amoyel et al., 2016b). EdU incorporation was carried out by incubating dissected samples in Schneider’s medium with 10μM EdU (Invitrogen) for 30 minutes prior to fixing. After primary and secondary antibody steps, samples were incubated in buffer containing 0.1mM THPTA, 2mM sodium ascorbate, 1mM CuSO_4_ and 2.5μM fluorescently-conjugated picolyl azide (Click Chemistry Tools) for 30 minutes, then briefly washed and mounted for imaging.

We used the following primary antibodies: rabbit anti-Zfh1 (1:5000, gift of R. Lehmann), guinea pig anti-Zfh1 (1:500, gift of J. Skeath), rabbit anti-phospho-4E-BP1 (1:200, Cell Signaling #2855), rabbit anti-phospho-Akt (1:200, Cell Signaling #9271), rabbit anti-phospho-S6k (1:200, Cell Signaling #9209), guinea pig anti-Tj (1:3000, gift of D. Godt), rabbit anti-GFP (1:500, Invitrogen #A-6455), chicken anti-GFP (1:500, Aves Labs #GFP-1010); rat anti-Vasa developed by D. Williams and A.C. Spradling (1:20), mouse anti-Fas3 (7G10) developed by C. Goodman (1:20), mouse anti-Eya (10H6), developed by S. Benzer and N.M. Bonini (1:20), rat anti-Ncad (DN-Ex#8), developed by T. Uemura (1:20), mouse anti-Dlg (4F3) developed by C. Goodman (1:20) were all obtained from the Developmental Studies Hybridoma Bank, created by the NICHD of the NIH and maintained at The University of Iowa, Department of Biology, Iowa City, IA 52242.

### Statistical analysis

Unless otherwise indicated, numbers shown are mean ± standard error. Significance for stem cell number counts was determined using Mann-Whitney tests for pairwise comparison, or nonparametric ANOVA (Kruskal-Wallis test) followed by Dunn’s multiple comparisons test to assess differences between multiple samples. For clone recovery rates, Fisher’s exact test was carried out. All P values were determined using GraphPad Prism software.

## Acknowledgements

We are grateful to members of the Amoyel, Fernandes, Poole and Barrios labs for productive discussions, to Vilaiwan Fernandes, Ben Simons, Claudio Stern, Richard Poole and Barbara Conradt for critical comments on the manuscript. Many thanks to Salvador Herrera for suggesting the use of Gal80 flipout cassettes and to members of the fly community for stocks and antibodies. Work in the Amoyel lab is funded by an MRC CDA.

## Author contributions

ACY, K-HH and MA performed and analysed experiments. MA conceived the project, designed the experiments and wrote the manuscript. All authors contributed to the article and approved the submitted version.

## References

Amoyel, M., Sanny, J., Burel, M., and Bach, E.A. (2013). Hedgehog is required for CySC selfrenewal but does not contribute to the GSC niche in the Drosophila testis. Dev. 140, 56–65.

Amoyel, M., Simons, B.D., and Bach, E.A. (2014). Neutral competition of stem cells is skewed by proliferative changes downstream of Hh and Hpo. EMBO J. 33, 2295–2313.

Amoyel, M., Hillion, K.H., Margolis, S.R., and Bach, E.A. (2016a). Somatic stem cell differentiation is regulated by PI3K/Tor signaling in response to local cues. Dev. 143, 3914–3925.

Amoyel, M., Anderson, J., Suisse, A., Glasner, J., and Bach, E.A. (2016b). Socs36E Controls Niche Competition by Repressing MAPK Signaling in the Drosophila Testis. PLoS Genet. 12.

Bohm, R.A., Welch, W.P., Goodnight, L.K., Cox, L.W., Henry, L.G., Gunter, T.C., Bao, H., and Zhang, B. (2010). A genetic mosaic approach for neural circuit mapping in Drosophila. Proc. Natl. Acad. Sci. U. S. A. 107, 16378–16383.

Brawley, C., and Matunis, E. (2004). Regeneration of male germline stem cells by spermatogonial dedifferentiation in vivo. Science (80-.). 304, 1331–1334.

Chatzeli, L., and Simons, B.D. (2020). Tracing the dynamics of stem cell fate. Cold Spring Harb. Perspect. Biol. 12.

Chen, D., and McKearin, D. (2003). Dpp Signaling Silences bam Transcription Directly to Establish Asymmetric Divisions of Germline Stem Cells. Curr. Biol. 13, 1786–1791.

Cheng, J., Tiyaboonchai, A., Yamashita, Y.M., and Hunt, A.J. (2011). Asymmetric division of cyst stem cells in Drosophila testis is ensured by anaphase spindle repositioning. Development 138, 831–837.

Chiang, A.C.Y., Yang, H., and Yamashita, Y.M. (2017). spict, a cyst cell-specific gene, regulates starvation-induced spermatogonial cell death in the Drosophila testis. Sci. Rep. 7.

Clevers, H., and Watt, F.M. (2018). Defining Adult Stem Cells by Function, not by Phenotype. Annu. Rev. Biochem. 87, 1015–1027.

Donati, G., and Watt, F.M. (2015). Stem cell heterogeneity and plasticity in epithelia. Cell Stem Cell 16, 465–476.

Enver, T., Pera, M., Peterson, C., and Andrews, P.W. (2009). Stem Cell States, Fates, and the Rules of Attraction. Cell Stem Cell 4, 387–397.

Fabrizio, J.J., Boyle, M., and DiNardo, S. (2003). A somatic role for eyes absent (eya) and sine oculis (so) in Drosophila spermatocyte development. Dev. Biol. 258, 117–128.

Fairchild, M.J., Smendziuk, C.M., and Tanentzapf, G. (2015). A somatic permeability barrier around the germline is essential for Drosophila spermatogenesis. Development 142, 268–281.

Ge, Y., and Fuchs, E. (2018). Stretching the limits: From homeostasis to stem cell plasticity in wound healing and cancer. Nat. Rev. Genet. 19, 311–325.

Gonczy, P., and DiNardo, S. (1996). The germ line regulates somatic cyst cell proliferation and fate during Drosophila spermatogenesis. Development 122.

Greenspan, L.J., de Cuevas, M., and Matunis, E. (2015). Genetics of Gonadal Stem Cell Renewal. Annu. Rev. Cell Dev. Biol. 31, 291–315.

Hara, K., Nakagawa, T., Enomoto, H., Suzuki, M., Yamamoto, M., Simons, B.D., and Yoshida, S. (2014). Mouse spermatogenic stem cells continually interconvert between equipotent singly isolated and syncytial states. Cell Stem Cell 14, 658–672.

Haramis, A.P.G., Begthel, H., Van Den Born, M., Van Es, J., Jonkheer, S., Offerhaus, G.J.A., and Clevers, H. (2004). De Novo Crypt Formation and Juvenile Polyposis on BMP Inhibition in Mouse Intestine. Science (80-.). 303, 1684–1686.

Hardy, R.W., Tokuyasu, K.T., Lindsley, D.L., and Garavito, M. (1979). The germinal proliferation center in the testis of Drosophila melanogaster. J. Ultrasructure Res. 69, 180–190.

He, X.C., Zhang, J., Tong, W.G., Tawfik, O., Ross, J., Scoville, D.H., Tian, Q., Zeng, X., He, X., Wiedemann, L.M., et al. (2004). BMP signaling inhibits intestinal stem cell self-renewal through suppression of Wnt-β-catenin signaling. Nat. Genet. 36, 1117–1121.

Jones, D.L., and Wagers, A.J. (2008). No place like home: Anatomy and function of the stem cell niche. Nat. Rev. Mol. Cell Biol. 9, 11–21.

Kawase, E., Wong, M.D., Ding, B.C., and Xie, T. (2004). Gbb/Bmp signaling is essential for maintaining germline stem cells and for repressing bam transcription in the Drosophila testis. Development 131, 1365–1375.

Kiger, A.A., White-Cooper, H., and Fuller, M.T. (2000). Somatic support cells restrict germline stem cell self-renewal and promote differentiation. Nature 407, 750–754.

Kirilly, D., Wang, S., and Xie, T. (2011). Self-maintained escort cells form a germline stem cell differentiation niche. Development 138, 5087–5097.

Klein, A.M., and Simons, B.D. (2011). Universal patterns of stem cell fate in cycling adult tissues. Development 138, 3103–3111.

Krieger, T., and Simons, B.D. (2015). Dynamic stem cell heterogeneity. Dev. 142, 1396–1406.

Leatherman, J.L., and Dinardo, S. (2010). Germline self-renewal requires cyst stem cells and stat regulates niche adhesion in Drosophila testes. Nat. Cell Biol. 12, 806–811.

Leatherman, J.L., and Di Nardo, S. (2008). Zfh-1 controls somatic stem cell self-renewal in the Drosophila testis and nonautonomously influences germline stem cell self-renewal. Cell Stem Cell 3, 44–54.

Lee, T., and Luo, L. (1999). Mosaic analysis with a repressible neurotechnique cell marker for studies of gene function in neuronal morphogenesis. Neuron 22, 451–461.

Lenhart, K.F., and DiNardo, S. (2015). Somatic Cell Encystment Promotes Abscission in Germline Stem Cells following a Regulated Block in Cytokinesis. Dev. Cell 34, 192–205.

Matunis, E., Tran, J., Gönczy, P., Caldwell, K., and DiNardo, S. (1997). Punt and schnurri regulate a somatically derived signal that restricts proliferation of committed progenitors in the germline. Development 124, 4383–4391.

Michel, M., Kupinski, A.P., Raabe, I., and Bökel, C. (2012). Hh signalling is essential for somatic stem cell maintenance in the drosophila testis niche. J. Cell Sci. 125, 2663–2669.

Morrison, S.J., and Spradling, A.C. (2008). Stem Cells and Niches: Mechanisms That Promote Stem Cell Maintenance throughout Life. Cell 132, 598–611.

Murayama, K., Kimura, T., Tarutani, M., Tomooka, M., Hayashi, R., Okabe, M., Nishida, K., Itami, S., Katayama, I., and Nakano, T. (2007). Akt activation induces epidermal hyperplasia and proliferation of epidermal progenitors. Oncogene 26, 4882–4888.

Nakagawa, T., Sharma, M., Nabeshima, Y.I., Braun, R.E., and Yoshida, S. (2010). Functional hierarchy and reversibility within the murine spermatogenic stem cell compartment. Science (80-.). 328, 62–67.

Schofield, R. (1978). The relationship between the spleen colony-forming cell and the haemopoietic stem cell. Blood Cells 4, 7–25.

Sênos Demarco, R., Uyemura, B.S., and Jones, D.L. (2020). EGFR Signaling Stimulates Autophagy to Regulate Stem Cell Maintenance and Lipid Homeostasis in the Drosophila Testis. Cell Rep. 30, 1101–1116.e5.

Sheng, X.R., Brawley, C.M., and Matunis, E.L. (2009). Dedifferentiating Spermatogonia Outcompete Somatic Stem Cells for Niche Occupancy in the Drosophila Testis. Cell Stem Cell 5, 191–203.

Shields, A.R., Spence, A.C., Yamashita, Y.M., Davies, E.L., and Fuller, M.T. (2014). The actin-binding protein profilin is required for germline stem cell maintenance and germ cell enclosure by somatic cyst cells. Dev. 141, 73–82.

Shivdasani, A.A., and Ingham, P.W. (2003). Regulation of Stem Cell Maintenance and Transit Amplifying Cell Proliferation by TGF-β Signaling in Drosophila Spermatogenesis. Curr. Biol. 13, 2065–2072.

Singh, S.R., Liu, Y., Zhao, J., Zeng, X., and Hou, S.X. (2016). The novel tumour suppressor Madm regulates stem cell competition in the Drosophila testis. Nat. Commun. 7.

Smendziuk, C.M., Messenberg, A., Wayne Vogl, A., and Tanentzapf, G. (2015). Bi-directional gap junction-mediated soma-germline communication is essential for spermatogenesis. Dev. 142, 2598–2609.

Stine, R.R., Greenspan, L.J., Ramachandran, K. V., and Matunis, E.L. (2014). Coordinate Regulation of Stem Cell Competition by Slit-Robo and JAK-STAT Signaling in the Drosophila Testis. PLoS Genet. 10.

Tai, K., Cockburn, K., and Greco, V. (2019). Flexibility sustains epithelial tissue homeostasis. Curr. Opin. Cell Biol. 60, 84–91.

Tamirisa, S., Papagiannouli, F., Rempel, E., Ermakova, O., Trost, N., Zhou, J., Mundorf, J., Brunel, S., Ruhland, N., Boutros, M., et al. (2018). Decoding the Regulatory Logic of the Drosophila Male Stem Cell System. Cell Rep. 24, 3072–3086.

Tran, J., Brenner, T.J., and DiNardo, S. (2000). Somatic control over the germline stem cell lineage during Drosophila spermatogenesis. Nature 407, 754–757.

Wang, S., Gao, Y., Song, X., Ma, X., Zhu, X., Mao, Y., Yang, Z., Ni, J., Li, H., Malanowski, K.E., et al. (2015). Wnt signaling-mediated redox regulation maintains the germ line stem cell differentiation niche. Elife 4.

Xie, T., and Spradling, A.C. (1998). decapentaplegic is essential for the maintenance and division of germline stem cells in the Drosophila ovary. Cell 94, 251–260.

Xie, T., and Spradling, A.C. (2000). A niche maintaining germ line stem cells in the Drosophila ovary. Science (80-.). 290, 328–330.

Ying, Q.L., Wray, J., Nichols, J., Batlle-Morera, L., Doble, B., Woodgett, J., Cohen, P., and Smith, A. (2008). The ground state of embryonic stem cell self-renewal. Nature 453, 519–523.

Ying, Z., Sandoval, M., and Beronja, S. (2018). Oncogenic activation of PI3K induces progenitor cell differentiation to suppress epidermal growth. Nat. Cell Biol. 20, 1256–1266.

Yoshida, S. (2018). Open niche regulation of mouse spermatogenic stem cells. Dev. Growth Differ. 60, 542–552.

